# Mutations alter RNA-mediated conversion of human prions

**DOI:** 10.1101/235879

**Authors:** Erik J. Alred, Izra D. Lodangco, Jennifer Gallaher, Ulrich Hansmann

## Abstract

Prion diseases are connected with self-replication and self-propagation of mis-folded proteins. The rate-limiting factor is the formation of the initial seed. We have recently studied early stages in the conversion between functional PrP^C^ and the infectious scrapie PrP^SC^ form, triggered by the binding of RNA. Here, we study how this process is modulated by the prion sequence. We focus on residues 129 and 178, which are connected to the hereditary neurodegenerative disease Fatal Familial Insomnia.

## Introduction

Prions play a critical role in the maintenance and growth of neuronal synapses in mammalian and avian species,^(1-3)^ and improperly folded prions are associated with various neurodegenerative diseases. Examples are scrapie in sheep, Bovine Spongiform Encephalopathy in cows, Chronic Wasting Disease in elks and deer; and Creutzfeldt-Jacob, Kuru, Fatal Familial Insomnia, and potentially even the pregnancy-specific disorder preeclampsia, in humans.^(3-6)^ The shared characteristic in all these illnesses are cytotoxic aggregates formed from mis-folded prions.

However, the underlying disease mechanism is difficult to probe as, unlike to the functional form PrP^C^, no structural models of mis-folded and toxic PrP^SC^ have been resolved and published in the Protein Data Bank (PDB). Circular dichroism measurements show for this so-called scrapie form PrP^SC^ a reduction in helicity from 43% in PrP^C^ to 30% in PrP^SC^, coupled with a rise in β-strand frequency of 3% in PrP^c^ to 43% in PrP^sc^. ^(7-14)^ For the mouse prion it is known that it is primarily the N-terminal helix A (residues 143–161) which decays, while the central helix B (172–196) and the C-terminal helix C (200–229) stay intact or dissolve later.^(7-14)^ It is also known that the PrP^SC^ structure is self-propagating, i.e., a prion in this form can convert functional PrP^C^ prions into its toxic PrP^SC^ form. The rate-limiting factor in this “protein-only” mechanism of transmission^(15)^ is the formation of the initial seed. We speculate that mutations which are known to lead to earlier outbreak, faster disease progression and more sever symptoms, cause these effects though enhancing the generation of the initial seeds. If true it would be therefore important for evaluating treatment options to understand the mechanism by that the seeds are generated, and how this mechanism is altered by mutations.

Various experimental^1, 16^ and computational^(17)^ studies point to the possibility that poly-adenosine RNA (poly-A-RNA) catalyzes conversion of the native PrP^C^ form into PrP^SC^ through interacting with the N-terminal of the prion. Simulating mouse and human prions, we have observed that binding of a poly-A-RNA fragment with either a polybasic domain of residues 21-31, or with a segment made form residues 144-155, leads to formation of a characteristic pincer motif between the polybasic domains of residues 21-31 and on helix A. This pincer encapsulates then RNA fragment, and formation of the resulting complex leads to unfolding of helix A as interactions between the RNA fragment and the side chains of residues 144-148 disrupt locally the backbone hydrogen bonds of helix A.^17,18^

In the present project we study whether and how this mechanism changes when going from wild type prions to mutants associated with sever illnesses. Fatal Familial Insomnia is a neurodegenerative disorder caused by loss of neurons in the thalamus^(19)^ that starts with progressively worsening insomnia which rapidly degenerates into dementia.^(20)^ There is currently no effective treatment, and patients die within 12 months of the first appearance of symptoms.^(21)^ The disease is associated with a heritable mutation D178N, replacing an acidic aspartic acid (D) by a neutral asparagine (N), which is known to increase the polymorphism of the scrapie form.^(22)^ The disease also requires a methionine (M) at position 129 and is not observed with a valine (V) at position 129, another often found variant of the wild type.^(23)^ In our previous investigation^(17)^ we considered only the predominant 129M variant of the wild type. Here, we extend these investigations and research how the residue replacements D178N and M129V alter the interactions between poly-A-RNA and prions and the mechanism that leads to unfolding of helix A. Four systems will be studied: the wild type with 129M and 178D (129M-178D), its variant with 129V (129V-178D), and the two mutants 129M-178N and 129V-178N. Analyzing long-time molecular dynamics simulations of these four systems we argue that residue 129 controls the probability of RNA binding to a site that allows for the pincer conversion mechanism proposed by us in earlier work,^(17)^ while the mutation D178N increases the stability and extension of the pincer.

## Results and Discussion

In Ref. 17 we have identified three sites where the poly-A-RNA can bind to the PrP^C^ form of the wild type 129M-178D. At binding site 1, the RNA interacts with residues 21-31, at binding site 2 with residues 111-121; and binding site 3 consists of residues 144 to 155. In the eleven stable complexes predicted by Autodock, binding site 1 was observed five times, site 2 four times, and site 3 three times. Only when the RNA binds to site 1 and site 3 did we see in molecular dynamic simulations the formation of a pincer motif followed by decay of the N-terminal helix A, the first step in the conversion to the scrapie form PrP^SC^. On the other hand, binding to site 2 did not alter the stability of helix A or any other helix.

The residue replacements M129V and/or D178N shift the probabilities of the binding sites but we do no find new binding sites in the three prions 129V-178D, 129M-178N, and129V-178N considered now also by us. For the 129M-178N mutant, binding site 3 is observed in all of the ten top complexes predicted by Autodock. For the 129V-178D sequential polymorph of the wild type, eight of the top ten complexes bind at site 2, with the remaining two complexes binding to site 1. For the last prion, the mutant 129V-178N, we again observe all three of our previously identified binding sites: site 3 is found six times, site 1 three times, and site 2 is seen once. As in our previous study of the 129M-178D wild type, we selected for all of the three new systems (129V-178D, 129M-178N and 129V-178N) the most probable conformation (see methods section for further explanation) of a complex with a certain binding site, and followed its time evolution over 300ns in three independent trajectories. A matrix of the various systems, binding sites and trajectory numbers is given in **Table 1**.

**Table 1:** Frequency of binding sites found in the Top 10 of Autodock predictions. For a given system and binding site the most stable complex is followed in three trajectories over 300 ns.

| System | Site 1 | Site 2 | Site 3 | Trajectories |
| --- | --- | --- | --- | --- |
| **129M-178D** | 4 | 3 | 3 | 3 x 3 |
| **129V-178D** | 2 | 8 | 0 | 3 x 2 |
| **129M-178N** | 0 | 0 | 10 | 3 x 1 |
| **129V-178N** | 1 | 3 | 6 | 3 x 3 |

## Visual Inspection and trajectory analysis

Fatal Familial Insomnia is connected with a methionine at residue 129 (129M) and the mutation D178N. Poly-A-RNA binding to this mutant (our system 129M-178N) is only observed at binding site 3, residues 144-155. In the wild type (129M-178D) studied in our previous work, binding of RNA to site 3 always lead to the unfolding of helix A. The same is observed here for the mutant; however, we find two differences. First, in about 70% of all configurations sampled in our molecular dynamics trajectories, residue 178N forms a hydrogen bond with 18D that re-strains the movement of helix C and keeps it in close proximity to both helix A and the polybasic domain of residues 21-31, see **Figure 1-A**. This, secondly, allows for the RNA to interact with both helix A (140-158) and helix C (200-220), see **Table 2**. As a consequence, not only helix A, but also most of helix C has dissolved in the final configuration of the 300ns trajectory. Note that helix C is now also participating in the three pronged helix-polybasic pincer motif, adding to the polybasic domain of residues 21-31 and the segment of residues 140-161 on helix A additional contacts between the RNA and residues 219-223 on helix C (see **Table 2**). However, while all trajectories led to unfolding of helix A, this enlarged pincer and the decay of helix C is only seen in two of the three trajectories.

**Figure 1:**
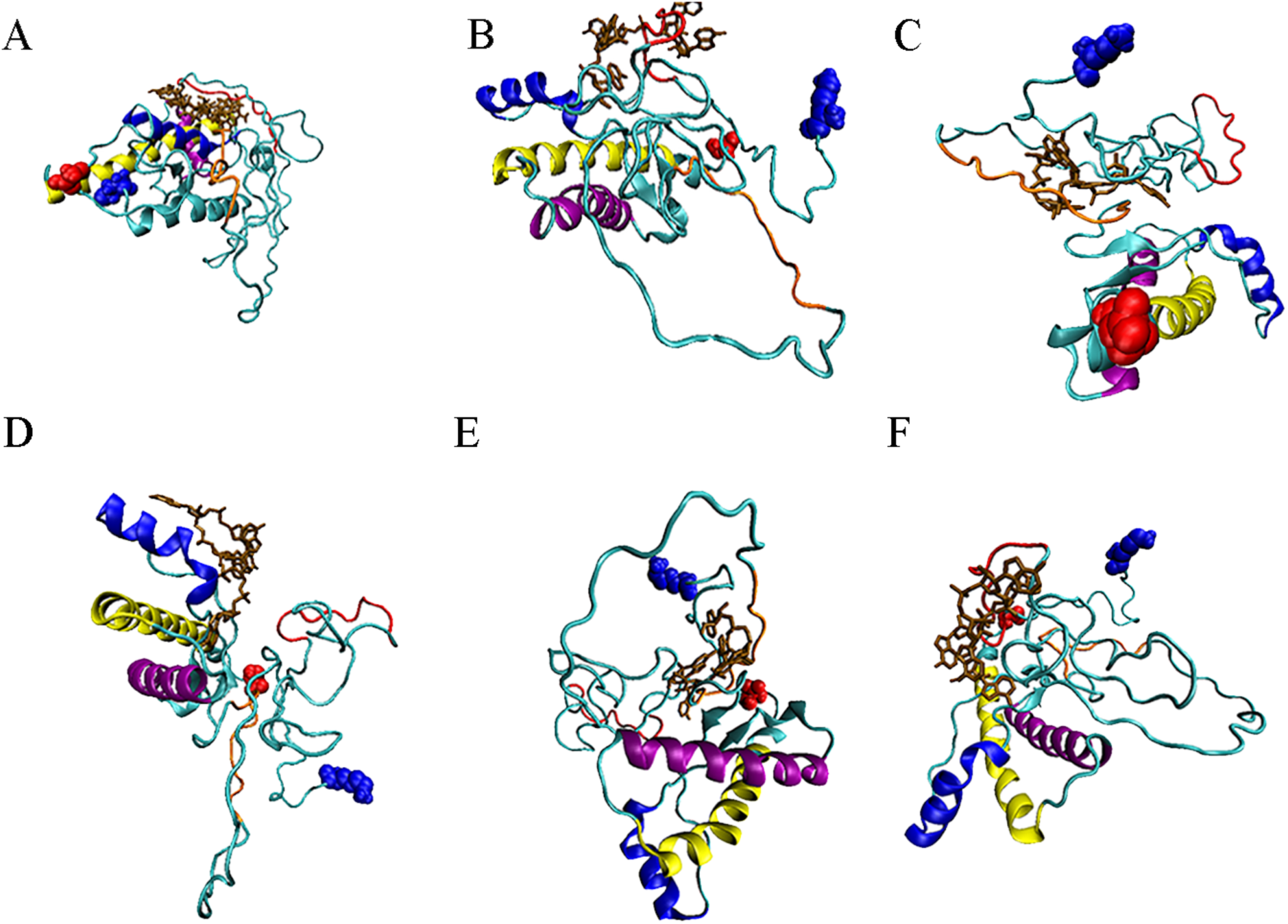
Initial configurations of complexes of RNA bound to our three prion models. The 129M-178N mutant binds only to site 3 shown in (A), while the 129V-178N binds to site 1 (B), site 2 (C), or site 3 (D). The 129V-178D wild type variant binds to either site 2 (E) or site 1 (F). Colors denote the following: Brown: RNA molecule, Blue helix: helix A, Yellow helix: helix B, Purple helix: helix C, Red strand: binding site 1, Orange strand: binding site 2, Blue ball: N-terminus, Red ball: C-terminus.

**Table 2:**
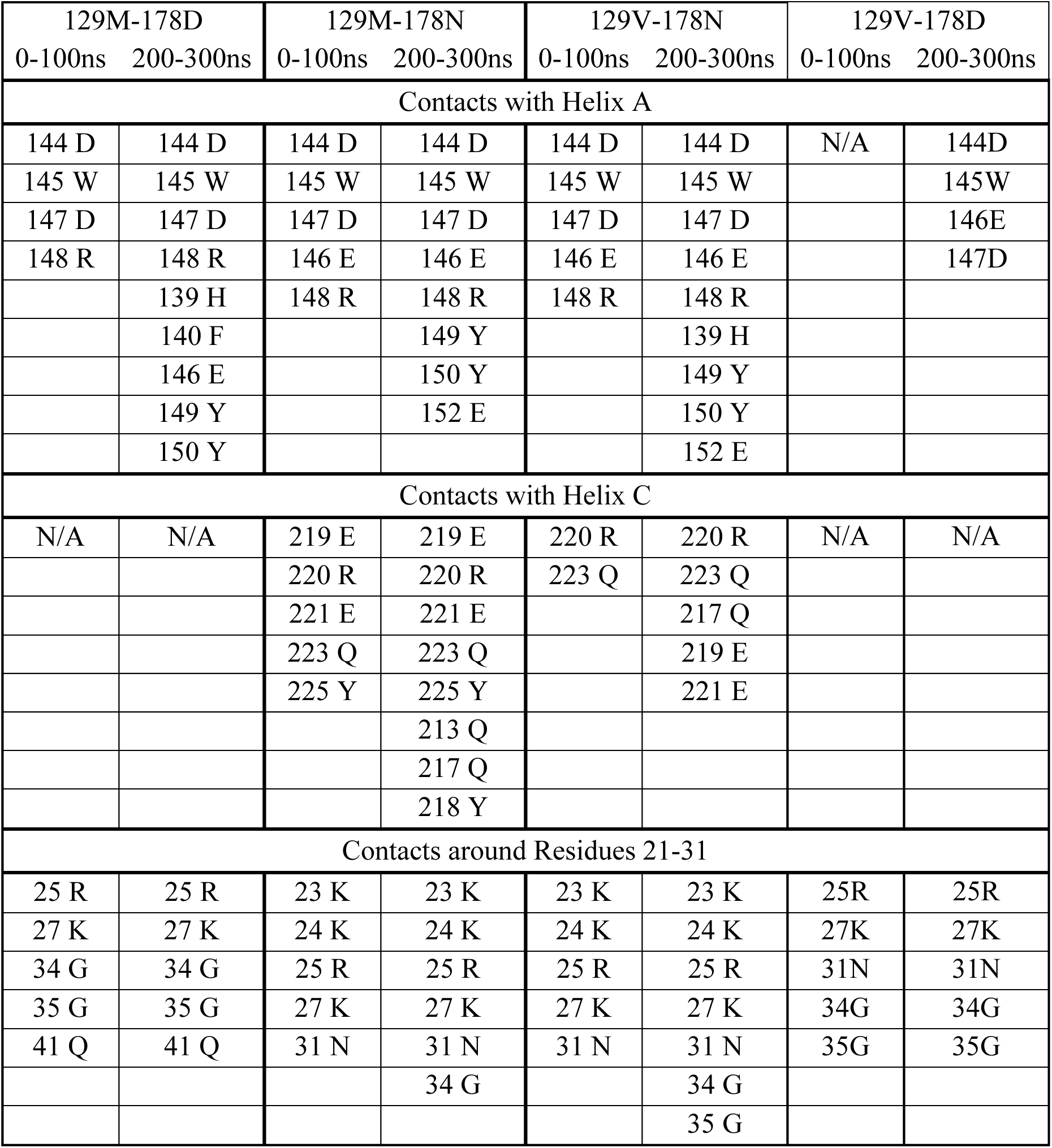
Contacts between the prion protein and the RNA-fragment that are formed in more than 50% of the time-steps. Data are for complexes where the RNA-fragment binds to either site 1 or site 3, leading to unfolding of helices.

The above picture is quantified in **Figure 2-B** where we show the time evolution of the root-mean-square-deviation (RMSD) with respect to the start configuration. We compare the complex of poly-A-RMA binding to the prion with that of the isolated prion having the same structure. For comparison, we show in **Figure 2-A** the same quantity as calculated from our previous studies of the wild type 129M-178D in Ref. 17. In all systems we distinguish between complexes where the RNA docks to binding site 1 or 3 (shown in red), and such where it docks to binding site 2 (shown in gold). The blue circle indicates the region of the trajectory after which less than 50% of the helical contacts of helix A remain. These are also the regions where we observe large changes in RMSD. This transition region appears much earlier in the trajectory for the mutant system. Note also the second jump in RMSD marked by the black circle where 50% of helical contacts for helix C have decayed. The decrease in helicity is also seen in **Table 3**. While the helicity does not change in the control, the average helicity decreases from about 43% to 32% for the wild type when RNA binds to either site 1 or 3. In the 129M-178N mutant the overall helicity decreases from 49% o 30%, most of it coming from helix A (94% o 18%), with the reduction for helix C from 96% to 39%.

**Figure 2:**
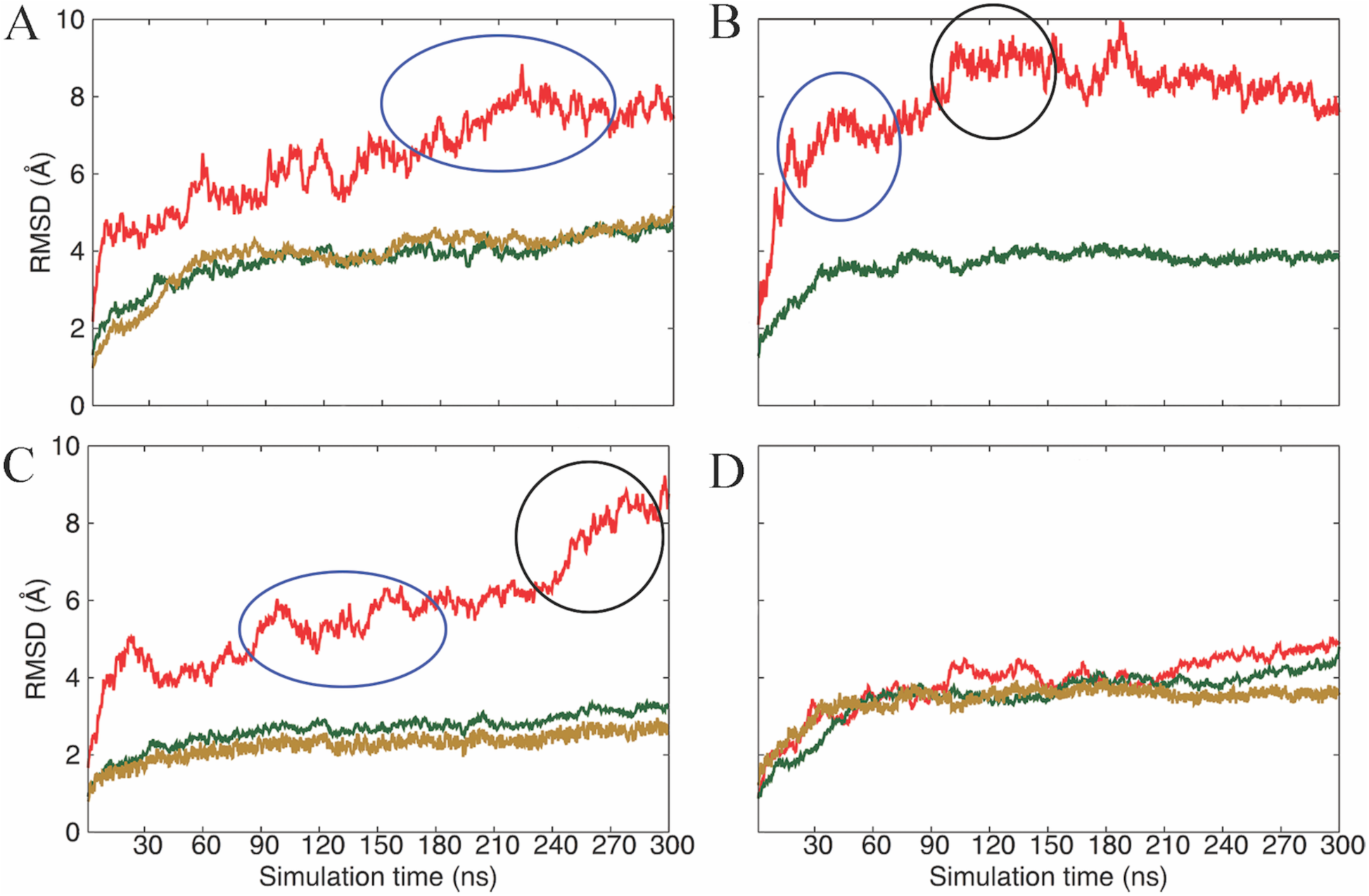
Root-mean-square-deviation (RMSD) of a configuration at time t to the start configuration for (A) wild type 129M-178D, mutant (B) 129M-178N and (C) 129V-178N, and (D) wild type variant 129V-178D. Shown are for each system averages over three trajectories. RMSD values for isolated prions are drawn in green, that for systems where the RNA binds to site 1 or 3 in red; and for systems where the RNA binds to site 2 in gold. Blue circles indicate timeframes where the chance of finding a contact with helix becomes less than 50%. The black circle marks the same point for helix C.

**Table 3:** Secondary structure averaged over specific periods of the respective trajectories. Data are for complexes where the RNA-fragment binds to either site 1 or site 3.

| Secondary Structure | 129M-178D |  |  | 129V-178D |  |  |
| --- | --- | --- | --- | --- | --- | --- |
|  | Control 0-300ns | Complex 0-100ns | Complex 200-300ns | Control 0-300ns | Complex 0-100ns | Complex 200-300ns |
| Total Helicity | 43% (2%) | 39% (3%) | 32% (2%) | 44% (2%) | 43% (1%) | 44% (1%) |
| Helix A | 92% (2%) | 56% (6%) | 22% (4%) | 97% (1%) | 97% (1%) | 97% (2%) |
| Helix C | 96% (2%) | 93% (3%) | 92% (3%) | 98% (2%) | 94% (1%) | 95% (2%) |
| β-Strands | 4% (1%) | 7% (2%) | 9% (2%) | 5% (1%) | 4% (1%) | 4% (1%) |
| β-Strand Occupancy | 28% (1%) | 38% (5%) | 58% (4%) | 28% (2%) | 25% (1%) | 27% (1%) |
| Secondary Structure | 129M-178N |  |  | 129V-178N |  |  |
|  | Control 0-300ns | Complex 0-100ns | Complex 200-300ns | Control 0-300ns | Complex 0-100ns | Complex 200-300ns |
| Total Helicity | 48% (2%) | 35% (2%) | 30% (3%) | 48% (3%) | 40% (3%) | 36% (5%) |
| Helix A | 96% (3%) | 51% (6%) | 18% (7%) | 98% (2%) | 60% (7%) | 25% (5%) |
| Helix C | 96% (3%) | 73% (3%) | 39% (5%) | 97% (3%) | 81% (4%) | 48% (5%) |
| β-Strands | 4% (1%) | 7% (2%) | 9% (2%) | 4% (2%) | 6% (2%) | 8% (3%) |
| β-Strand Occupancy | 26% (1%) | 41% (3%) | 60% (4%) | 27% (2%) | 40% (4%) | 52% (5%) |

Fatal Familial Insomnia is not observed in humans carrying the D178N mutation if residue 129 is a valine instead of the at this position more commonly observed methionine. Taking the 129M-178N mutant and substituting a valine at position 129, the 129V-178N mutant leads to poly-A-RNA binding to all three binding sites that are also observed in our previous simulations^(17)^ of the wild type 129M-178D. Complexes with these three binding sites are shown in **Figure 1 B-D**. Similar to the behavior of the 129M-178D wild type, unraveling of helices for the mutant 129V-178N is observed only for binding at sites 1 or 3, not when the poly-A-RNA fragment binds to site 2. However, **Table 2** also shows that if the pincer motif is formed, it is three-pronged as observed in the 129M-178N mutant, and involves helix A, helix C and the polybasic domain of residues 21-31. This motif is observed 3 in two of three trajectories for both binding site 1 and binding site.

The above observations are again quantified by the RMSD plots in **Figure 2-C**. Initially, the docked system resembles the 129M-178D wild type (**Figure 2-A**), with similar regions for the decays of helix A (indicated by blue circles). At 240ns we also see a jump in RMSD corresponding to the unraveling of helix C indicated by a black circle, however, the signal is less pronounced than for the 129M-178N mutant. Correspondingly, the decline in helicity in **Table 3** is less than see for the 129M-178N mutant, decreasing from 97% to 25% in helix A, and 97% to 48% in helix C. Hence, comparing the two mutants 129M-178N and 129V-178N it appears that the mutation D178N extends the unraveling of helices from helix A to helix C. On the other hand, a valine instead of a methionine as residue 129 seems to increase the frequency of binding to site 2 which does not lead to unraveling of helices in the prion protein.

In the wild type, we observe transient β-strands in the region of helix A that hint at the start of the conversion to the PrP^SC^ state, as shown by the increase in relative β-strand content and occupancy in **Table 3**. Here, occupancy is defined as the average amount of time a β-strand is observed. While the total β-strand propensity increases only by about 5%, the average life time of the transient β-strands grows by an about 25%, which may indicate that the conversion to the β-sheet rich PrP^SC^ structure is to begin. Such transient β-strands are also observed in both D178N mutants, with similar values for β-strand content and occupancy. However, even with the more rapid and extensive helical unfolding seen in the D178N mutants, 300 ns is clearly too short for formation of the stable β-sheets expected in the PrP^SC^.

From our comparison of the two mutants 129M-178N and 129V-178N, we would expect to see for the wild type with valine at position 129 (129V-178D) that the frequency of complexes with poly-A-RNA binding to site 2 is larger than seen for the 129M-178D wild type. We find indeed that for 129V-178D in 80% of the complexes poly-A-RNA binds to site 2 (**Figure 1-E**), while the corresponding number in our previous simulations of 129M-178D is only 40%.^(17)^ As observed in all our simulations, binding to site 2 does not lead to unfolding of helices. Only in 20% of cases did we find binding to site 1 (**Figure 1-F**), and in no case binding to site 3. On the other hand, binding site 1 and 3 are observed with a frequency of 80% in the 129M-178D wild type. While complexes with binding site 1 are observed with lower frequency for the 129V-178D wild type than for the 129M-178D wild type, they lead again decay of helix A. However, the interaction is weaker, with only a partial unraveling of helix A that starts late after ~150ns and is not preceded by formation of the pincer motif. This can be seen from the RMSD plot in **Figure 2-D** where we do not see a discernable loss in helical contacts until the last 50ns of the trajectory. In addition, there is little difference between the RMSD values for binding site 1 (shown in red), binding site 2 (shown in gold) and the undocked system (shown in blue). Hence, it seems that valine instead of methionine at position 129 decreases not only the probability to bind to site 1 or site 3, but also in the rare cases where binding to these sites is seen, hinders or delays the formation of the pincer by increasing flexibility in critical regions of the N terminal domain.

## Hydrogen bond pattern and structural flexibility

In order to understand how the variation in sequence at position 129 and 178 changes the effect of poly-A-RNA binding to prions we focus in the following on trajectories where the RNA fragment binds to either site 1 or 3. This is because binding to site 2 never lead to unraveling of helices in the prion protein which we assume to be the initial state in the conversion from PrP^C^ to the scrapie form PrP^SC^. We start by looking in **Figure 3** into the root-mean-square-fluctuation (RMSF) of each residue in the various systems. The presented data are for complexes where the RNA fragment binds to the prion at either site 1 or 3, and are divided by the corresponding values for the unbound prion. Thus values greater than one indicate that the prion has grown more flexible upon binding with RNA. We show in **Figure 3-A** as reference our results from the previous study in Ref. 17 of the wild type 129M-178D. Note the characteristic spikes around the segment that forms helix A (residues 140-161) in the prion, indicating the unraveling of this helix upon binding of poly-A-RNA at site 1 or 3. This is confirmed by **Table 4** where the average probability of finding a 1-4 backbone hydrogen bonds in helix A is listed for a given time period. For comparison, we also show the corresponding frequencies for the undocked protein. Upon binding of the RNA fragment to either site 1 or 3 helix A loses contacts between residues 144-148 in the initial 100ns of the trajectory. As the simulations evolve, E146, Y49, Y150 and N153 lose their helical contacts as they begin to interact with the RNA. The time evolution of backbone-backbone contacts in this and the other three systems can be also seen from the contact maps shown in **Supplemental Figures S1-S3**.

**Figure 3:**
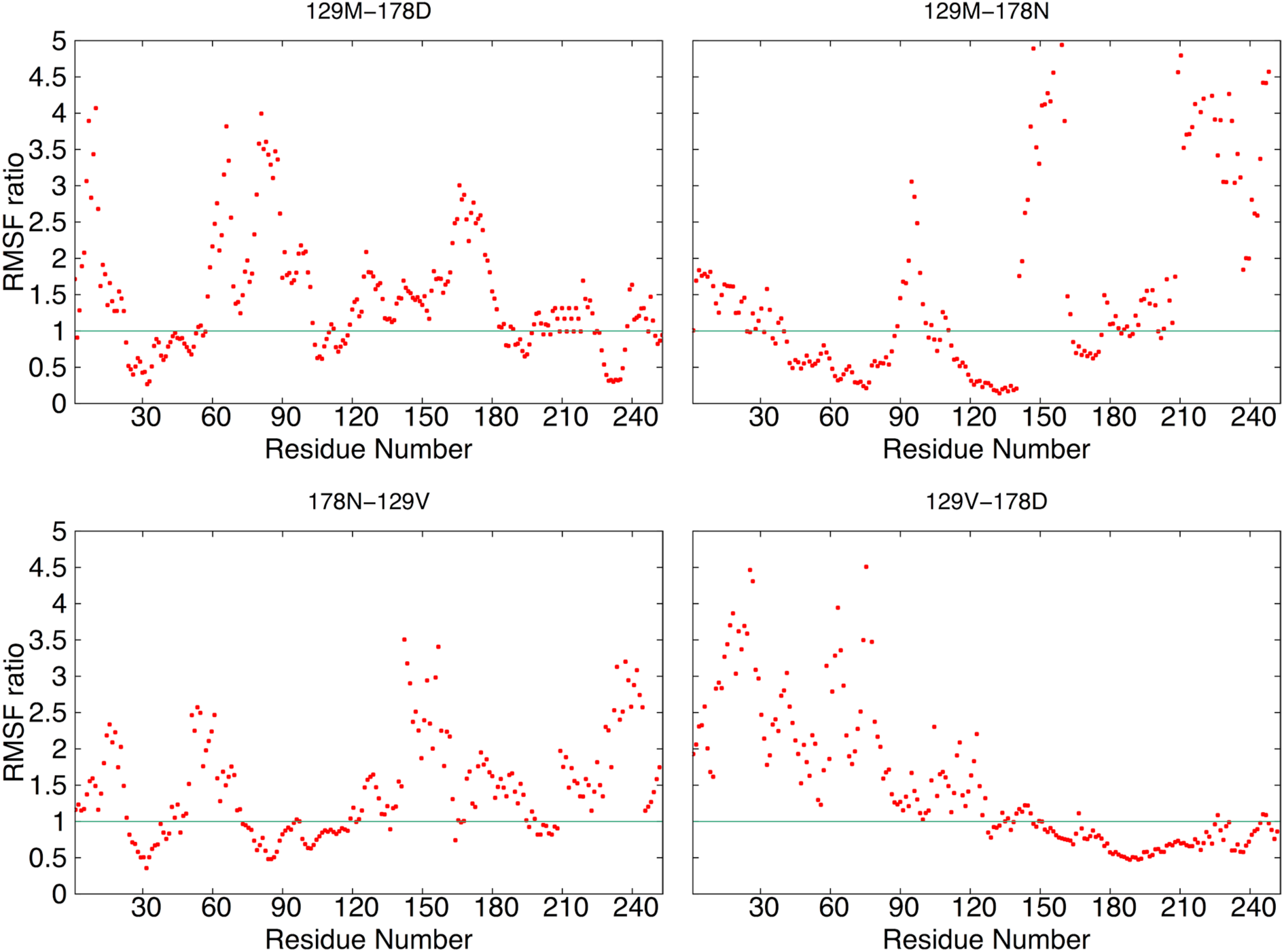
Relative root-mean-square fluctuation (RMSF) of residues. Shown is the ratio of RMSF values calculated for residues in prions where an RNA-fragment binds to either site 1 or site 2 (leading to unfolding of helices), divided by the corresponding values for the undocked prions. A ratio of one (marked by a green line) means that a given residue is equally flexible in the bound system and in the isolated prion. Values above the green line indicate elevated flexibility upon interaction with RNA.

**Table 4:** Helical contacts in helix A. Data are for complexes where the RNA-fragment binds to either site 1 or site 3.

|  | 129M-178D |  |  | 129V-178D |  |  |
| --- | --- | --- | --- | --- | --- | --- |
| Backbone<br>Hydrogen Bond | Control<br>0-300ns | Complex<br>0-100ns | Complex<br>200-300ns | Control<br>0-300ns | Complex<br>0-100ns | Complex<br>200-300ns |
| V161-P157 | 50% (4%) | 45% (5%) | 10% (3%) | 52% (3%) | 51% (3%) | 51% (4%) |
| V160-R156 | 75% (1%) | 55% (3%) | 16% (3%) | 83% (3%) | 78% (4%) | 76% (5%) |
| Q159-H155 | 89% (4%) | 62% (4%) | 20% (4%) | 90% (3%) | 88% (4%) | 80% (3%) |
| N158-M154 | 82% (4%) | 65% (4%) | 20% (4%) | 81% (3%) | 77% (4%) | 79% (4%) |
| P157-N153 | 46% (4%) | 43% (4%) | 20% (4%) | 47% (5%) | 48% (3%) | 53% (4%) |
| R156-E152 | 79% (2%) | 66% (8%) | 21% (4%) | 82% (4%) | 77% (5%) | 79% (6%) |
| H155-R151 | 98% (4%) | 81% (8%) | 51% (3%) | 96% (4%) | 94% (7%) | 86% (6%) |
| M154-Y150 | 74% (3%) | 66% (7%) | 31% (4%) | 77% (4%) | 79% (6%) | 71% (5%) |
| N153-Y149 | 86% (4%) | 50% (4%) | 0% (0%) | 89% (4%) | 86% (5%) | 92% (4%) |
| E152-R148 | 97% (2%) | 72% (9%) | 22% (4%) | 96% (4%) | 92% (6%) | 89% (5%) |
| R151-D147 | 95% (3%) | 41% (5%) | 21% (4%) | 93% (5%) | 94% (5%) | 84% (5%) |
| Y150-E146 | 99% (1%) | 60% (5%) | 10% (2%) | 95% (4%) | 95% (6%) | 85% (6%) |
| Y149-W145 | 92% (5%) | 41% (4%) | 15% (5%) | 92% (4%) | 85% (4%) | 82% (4%) |
| R148-D144 | 76% (5%) | 43% (6%) | 23% (5%) | 77% (3%) | 66% (6%) | 65% (4%) |
| D147-R143 | 97% (2%) | 62% (5%) | 26% (3%) | 96% (4%) | 95% (3%) | 91% (4%) |
|  | 129M-178N |  |  | 129V-178N |  |  |
| V161-P157 | 50% (3%) | 50% (3%) | 16% (4%) | 53% (4%) | 53% (4%) | 14% (3%) |
| V160-R156 | 83% (2%) | 54% (4%) | 12% (6%) | 74% (4%) | 62% (4%) | 12% (6%) |
| Q159-H155 | 88% (5%) | 64% (4%) | 26% (6%) | 88% (4%) | 68% (5%) | 22% (7%) |
| N158-M154 | 84% (4%) | 62% (4%) | 19% (6%) | 81% (3%) | 68% (4%) | 18% (4%) |
| P157-N153 | 48% (6%) | 44% (4%) | 15% (4%) | 47% (4%) | 23% (4%) | 16% (6%) |
| R156-E152 | 79% (4%) | 62% (6%) | 19% (6%) | 78% (4%) | 59% (8%) | 19% (5%) |
| H155-R151 | 96% (4%) | 72% (9%) | 34% (5%) | 97% (3%) | 74% (9%) | 29% (6%) |
| M154-Y150 | 72% (5%) | 78% (8%) | 23% (4%) | 70% (8%) | 53% (7%) | 18% (4%) |
| N153-Y149 | 87% (3%) | 54% (6%) | 6% (6%) | 85% (5%) | 51% (9%) | 11% (6%) |
| E152-R148 | 96% (3%) | 71% (8%) | 15% (4%) | 95% (4%) | 65% (8%) | 16% (4%) |
| R151-D147 | 95% (3%) | 44% (5%) | 11% (6%) | 96% (5%) | 61% (7%) | 14% (6%) |
| Y150-E146 | 93% (6%) | 64% (6%) | 8% (6%) | 94% (3%) | 65% (7%) | 14% (7%) |
| Y149-W145 | 93% (2%) | 45% (5%) | 17% (6%) | 95% (5%) | 69% (6%) | 16% (6%) |
| R148-D144 | 78% (3%) | 40% (6%) | 11% (5%) | 80% (4%) | 62% (7%) | 10% (5%) |
| D147-R143 | 98% (2%) | 53% (5%) | 16% (6%) | 94% (6%) | 64% (8%) | 12% (7%) |

The spike in RMSF is larger in the mutant 129M-178N than in the wild type and leads to a similar reduction of 1-4 hydrogen bonds in helix A (**Table 4**). However, a second spike in RMSD is observed at residues 200-220, signaling the decreased stability of helix C, see **Figure 3-B** and **Table 5**. The higher flexibility in these two regions is, as seen in the wild type, associated with a drop in flexibility for the N-terminal residues 20-45 upon the formation of the helical-polybasic pincer motif.

**Table 5:** Helical contacts for helix C. Data are for complexes where the RNA-fragment binds to either site 1 or site 3.

|  | 129M-178D |  |  | 129V-178D |  |  |
| --- | --- | --- | --- | --- | --- | --- |
| Backbone<br>Hydrogen Bond | Control<br>0 -300ns | Complex<br>0-100ns | Complex<br>200-300ns | Control<br>0-300ns | Complex<br>0-100ns | Complex<br>200-300ns |
| Q227-Q223 | 36% (5%) | 27% (6%) | 25% (7%) | 35% (5%) | 27% (6%) | 35% (5%) |
| Y226-S222 | 41% (5%) | 40% (4%) | 32% (4%) | 39% (5%) | 40% (5%) | 30% (5%) |
| Y225-E221 | 59% (5%) | 54% (3%) | 69% (4%) | 62% (7%) | 58% (7%) | 55% (6%) |
| A224-R220 | 66% (6%) | 58% (6%) | 71% (6%) | 67% (6%) | 69% (5%) | 76% (7%) |
| Q223-E219 | 78% (5%) | 79% (4%) | 73% (5%) | 74% (5%) | 69% (6%) | 65% (6%) |
| S222-Y218 | 91% (3%) | 89% (4%) | 96% (4%) | 89% (4%) | 88% (6%) | 90% (6%) |
| E221-Q217 | 94% (4%) | 94% (4%) | 93% (4%) | 92% (3%) | 81% (7%) | 97% (3%) |
| R220-T216 | 95% (3%) | 85% (5%) | 87% (5%) | 97% (3%) | 90% (5%) | 86% (5%) |
| E219-I215 | 94% (5%) | 85% (5%) | 87% (6%) | 92% (3%) | 92% (4%) | 90% (3%) |
| Y218-C214 | 89% (4%) | 78% (7%) | 98% (5%) | 90% (4%) | 87% (4%) | 94% (5%) |
| Q217-M213 | 88% (4%) | 85% (4%) | 87% (5%) | 92% (3%) | 88% (5%) | 91% (4%) |
| T216-Q212 | 88% (3%) | 81% (5%) | 93% (6%) | 95% (3%) | 87% (5%) | 91% (4%) |
| I215-E211 | 87% (5%) | 90% (4%) | 94% (5%) | 91% (3%) | 94% (4%) | 98% (5%) |
| C214-V210 | 79% (5%) | 72% (4%) | 81% (5%) | 79% (5%) | 66% (7%) | 75% (7%) |
| M213V209 | 95% (4%) | 89% (4%) | 91% (5%) | 90% (4%) | 91% (4%) | 97% (3%) |
| Q212-R208 | 92% (3%) | 90% (5%) | 98% (3%) | 93% (2%) | 84% (7%) | 96% (5%) |
| E211-E207 | 73% (6%) | 66% (6%) | 74% (5%) | 78% (6%) | 82% (7%) | 82% (6%) |
| V210-M206 | 88% (5%) | 80% (6%) | 85% (4%) | 94% (4%) | 83% (7%) | 86% (5%) |
| V209-M205 | 79% (5%) | 81% (5%) | 84% (5%) | 82% (3%) | 85% (6%) | 78% (7%) |
| R208-K204 | 92% (3%) | 85% (6%) | 86% (4%) | 95% (3%) | 86% (5%) | 81% (5%) |
| E207-V203 | 93% (5%) | 86% (4%) | 96% (4%) | 98% (3%) | 94% (3%) | 83% (5%) |
| M206-D202 | 97% (4%) | 88% (5%) | 87% (4%) | 92% (3%) | 90% (5%) | 91% (4%) |
| M205-T201 | 70% (5%) | 73% (5%) | 76% (5%) | 78% (5%) | 76% (3%) | 71% (7%) |
| K204-E200 | 91% (5%) | 87% (6%) | 83% (3%) | 88% (5%) | 82% (6%) | 93% (3%) |
|  | 129M-178N |  |  | 129V-178N |  |  |
| Q227-Q223 | 27% (5%) | 15% (7%) | 14% (5%) | 30% (5%) | 22% (4%) | 17% (6%) |
| Y226-S222 | 44% (5%) | 30% (7%) | 12% (6%) | 31% (5%) | 25% (5%) | 11% (7%) |
| Y225-E221 | 67% (8%) | 51% (9%) | 15% (4%) | 62% (7%) | 58% (3%) | 10% (8%) |
| A224-R220 | 59% (7%) | 40% (8%) | 11% (7%) | 69% (6%) | 69% (5%) | 21% (9%) |
| Q223-E219 | 74% (3%) | 42% (6%) | 19% (6%) | 79% (5%) | 67% (5%) | 17%(10%<br>) |
| S222-Y218 | 93% (2%) | 64% (6%) | 28% (5%) | 89% (4%) | 75% (6%) | 21% (8%) |
| E221-Q217 | 92% (3%) | 48% (7%) | 22% (7%) | 84% (6%) | 70% (6%) | 14% (8%) |
| R220-T216 | 95% (3%) | 63% (7%) | 29% (7%) | 94% (5%) | 85% (5%) | 33% (7%) |
| E219-I215 | 86% (4%) | 61% (8%) | 21% (5%) | 92% (3%) | 87% (4%) | 15% (6%) |
| Y218-C214 | 89% (5%) | 54% (9%) | 27% (6%) | 91% (4%) | 76% (5%) | 26% (8%) |
| Q217-M213 | 94% (3%) | 42% (9%) | 34% (6%) | 87% (4%) | 83% (4%) | 40% (6%) |
| T216-Q212 | 91% (4%) | 53% (8%) | 37% (6%) | 86% (4%) | 72% (4%) | 42% (7%) |
| I215-E211 | 89% (3%) | 49% (7%) | 35% (6%) | 87% (4%) | 76% (5%) | 42% (5%) |
| C214-V210 | 73% (6%) | 63% (6%) | 34% (7%) | 72% (6%) | 59% (4%) | 39% (6%) |
| M213V209 | 89% (4%) | 57% (8%) | 34% (6%) | 98% (3%) | 89% (5%) | 26% (7%) |
| Q212-R208 | 88% (4%) | 63% (5%) | 25% (5%) | 93% (5%) | 97% (5%) | 16% (8%) |
| E211-E207 | 82% (4%) | 79% (8%) | 29% (5%) | 81% (5%) | 77% (6%) | 20% (7%) |
| V210-M206 | 98% (2%) | 85% (7%) | 49% (5%) | 92% (4%) | 82% (5%) | 52% (7%) |
| V209-M205 | 87% (3%) | 76% (7%) | 38% (7%) | 77% (5%) | 72% (4%) | 31% (8%) |
| R208-K204 | 86% (5%) | 76% (7%) | 50% (6%) | 98% (5%) | 95% (4%) | 44% (7%) |
| E207-V203 | 97% (4%) | 86% (7%) | 87% (6%) | 93% (5%) | 95% (5%) | 93% (6%) |
| M206-D202 | 95% (4%) | 88% (6%) | 87% (3%) | 88% (4%) | 79% (4%) | 84% (7%) |
| M205-T201 | 70% (7%) | 71% (4%) | 72% (4%) | 72% (4%) | 60% (5%) | 69% (5%) |
| K204-E200 | 89% (6%) | 80% (4%) | 88% (4%) | 91% (4%) | 83% (4%) | 86% (8%) |

As expected, a similar but less pronounced behavior is also observed in the 129V-178N mutant (**Figure 3-C**), with the unraveling of helix C delayed (**Table 5**). Note especially, that binding of poly-A-RNA to the two mutants does not increase the flexibility of residues 45-90 as it does for the wild type 129M-178D. The increased flexibility of these residues in the later system results from a loss of contacts between helix A and polar residues of the N-terminal domain upon interaction with the RNA fragment. In the wild type, the frequency of these contacts drops from 32% (7%) for the isolated prion to 12% (3%) when in complex with RNA. However, in the two mutants there is the possibility for a contact between residue 178N and 18D that stabilizes the segment. As consequence, the frequency of contacts upon binding of RNA does not drop in the 129M-178N mutant, 39% (4%) versus 38% (7%). Hence, on effect of the D178N mutation seems to be the stabilization of the N-terminal segment of residues 45-90, which makes it easier to form the pincer that is associated with unraveling of the helices in the prion. This effect is weaker in the 129V-178N mutant where the frequency of stabilizing contacts still drops from 39% (6%) to 22% (4%).

On the other hand, the drop in frequency of these contacts in the 129V-178D variant of the wild type is similar to that in the 129M-178N form: 31% (4%) down to 11% (6%), however, the relative fluctuations in the N-terminal region are larger than in the 129M-178D wild type, see **Figure 3-D**. Hence, a valine instead of a methionine at position 129 may not only shift he binding pattern to site 2, but also increase the flexibility of the N-terminus. Since for 129V-178D the RNA fragment binds only to site 1 (residues 21-31) or 2 (residues 111-121), the higher flexibility of this region may explain the difficulties seen in forming a stable pincer motif and the late unraveling of helix A seen for this variant of the wild type.

**Table 6** shows that despite the loss of backbone hydrogen bonds in helix the newly formed contacts of residues with the poly-A-RNA result in an overall higher number of hydrogen bonds. For the Fatal Familial Insomnia causing sequence 129M-178N, 14 hydrogen bonds are gained despite the dissolution of helix A and C, possibly indicating the start of forming another ordered structure. For the wild type and the non-disease causing mutant 129V-178N the gain is still twelve hydrogen bonds. Hence, for the two mutants and the 129M wild type despite the loss of helix-stabilizing hydrogen bonds, the binding of RNA appears energetically favorable. This is different for the 129V variant of the wild type where due to the increased flexibility of the N-terminus about five hydrogen bonds are lost.

**Table 6:** Frequency of main-chain hydrogen bonds in complexes and the controls, and their difference Δ. Data are for complexes where the RNA-fragment binds to either site 1or site 3.

|  | 0-100ns |  |  | 200-300ns |  |  |
| --- | --- | --- | --- | --- | --- | --- |
| Name | Control | Complex | $\Delta$ | Control | Complex | $\Delta$ |
| **129M-178D** | 75 (3) | 82 (2) | 7 (4) | 82 (4) | 93 (3) | 11 (4) |
| **129V-178D** | 73 (4) | 75 (3) | 2 (5) | 80 (4) | 75 (4) | -5 (5) |
| **129M-178N** | 92 (3) | 99 (3) | 7 (4) | 93 (4) | 106 (3) | 13 (5) |
| **129V-178N** | 77 (3) | 81 (2) | 4 (4) | 75 (3) | 88 (3) | 12 (4) |

## Conclusion

We have extended a previous investigation^(17)^ of the effect of poly-A-RNA on the conversion of functional PrP^C^ into the toxic and self-propagating scrapie form PrP^SC^, by exploring how this process depends on the prion sequence. Our guiding assumption is that mutants which are associated with fast disease progression and sever symptoms ease the process of seed-generation. The example we chose is the D178N mutant which, when going together with a methione as residue 129, 129M-178N, leads to Fatal Familial Insomnia, but not if residue 129 is a valine (129V-178N). As controls we also looked into the case of the wild type variant 129V-178D (instead of the 129M-178D version studied by us in previous work^(17)^) which has a valine as residue 129 and is known to be associated with a lower frequency of neurodegenerative disorders. In a meta-analysis of Creutzfeldt-Jakob patients, only 10% were found to be homozygous for valine at the 129 position, with ~50% being homozygous for methionine at the 129 position, and the remaining being heterozygous. ^(24)^

In all cases we observe that unraveling of helix A is connected with the appearance of a pincer-like motif between helix A and the polybasic domain that encapsulating the RNA. Formation of the pincer requires binding of the RNA to either the polybasic segment of residues 21-31 (binding site 1) or the segment 141-155 (binding site 3), and traps the RNA. In both D178N mutants this pincer becomes three-pronged and leads also to unraveling of helix C. As the molecular dynamic trajectories proceed, formation of the pincer is followed by replacement of helical contacts in helix A (and helix C in the mutants) by short and transient β-strands that eventually may lead to the high β-sheet propensity of the disease-causing mis-folded PrP^SC^ structure. Hence, RNA binding to the prion can trigger the conversion of the cellular prion protein structure PrP^C^ to its infectious scrapie PrP^SC^ form, a process whose early steps (formation of the pincer motif starting unfolding of helix A and in the mutants helix C) could be observed in our 300 ns long molecular dynamics trajectories.

The sequence variation at position 129 and 178 of the prion protein modulate this process in two ways. First, they alter the probability that RNA binds to the prion protein at a site that allows formation of the pincer motif, and secondly, they change the stability and extension of this motif. The gatekeeper for the first effect appears to be residue 129, with a valine at this position skewing the binding of RNA to site 2, which does not lead to unraveling of helices, and further increasing the flexibility of the N-terminus. On the other hand, a methionine at this position increases the chances of binding to site 1 or site 3, resulting in formation of the pincer and subsequent unfolding of helix A. The second effect is controlled by residue 178. This mutation decreases the frequency of the side-chain contact between 178D-101K, which in turn increases the flexibility in residues 90-105 around binding site 2. Hence, the probability to bind to site 2 while at the same time binding to site 3 will be enhanced. An even more important effect of the mutation D178N is formation a stable contact between residues 178N and 18D that restricts the motion of the polybasic domain of residues 21-31, enhancing the appearance of the pincer motif. Furthermore, the 178N-18D contact brings the RNA into close proximity with the C terminal region of helix C, allowing for the formation of a three-pronged pincer between the polybasic domain and the 2 helices. This leads to the eventual decay of helical contacts in both helix A and C. The residue combination 129M and 178N both increases the frequency of binding of RNA to sites that allow for formation of the pincer motif, and the strength and extension of this motif. The net-effect is a faster conversion of the functional PrP^C^ prion structure to an infectious scrapie PrP^SC^ form that may seed formation of toxic amyloids which then cause the symptoms of Familial Fatal Insomnia.

## Materials and Methods

### Model Generation

In human systems, prion proteins attach to the cell membrane of neurons in the extracellular space via a glycosylphosphatidylinositol (GPI) anchor which is added to the C-terminus following a prior cleavage of 24 residues from the C-terminal residues.^(2-3)^ As our previous study showed that an unanchored system is sufficient to model the initial stages of conversion,^(17)^ we save on computational costs by modeling all our systems in this study with unanchored prions. In order to exclude erroneous charge interactions in the C-terminal region, we simulate the full-sized, non-cleaved, prions with all 253 residues. As only the C-terminal region of the native protein structure has been fully resolved, we have used in our previous work^(17)^ the two programs MODELLER and ITASS for prediction of the un-resolved N-terminal segment of residues 1-121^(25-29)^. Both methods led to similar models with marginal differences in binding sites and strength. As the ITASS structure preserved slightly the C-terminal helices, we used this protein structure as start point for our study. Using ITASS to refine our model to our previously determined template, we took this wild type Prp^C^ structure and mutated residues 178D and/or 129M into 178N and/or 129 V. This led to three structures, corresponding to 129M and 178N, 129V and 178D, 129V and 178N, which were relaxed in short (5 ns) molecular dynamics simulations at 310 K and 1 bar. Configurations were taken from these trajectories at 500 ps intervals (0.5 ns) and assessed using three separate methods for quantifying model quality: RAMPAGE ^(30)^, ERRAT^(31)^ and ProQ^(32)^. For this purpose, we calculate the the average score for the corresponding trajectory. If this score is below the cut-off value for any of the three methods, the initial structure is refined using a combination of MODELLER and ITASS as outlined in Ref. ^(33)^ until the resulting trajectory passes the quality threshold. The so-generated three structures were selected for the next stage of model generation.

### Docking Confirmation

As in our previous work^(17)^ we selected Autodock 25 to generate our protein-RNA complexes, a program that has a successful history of modeling similar systems.^(34-35)^ A 5-nucleotide snippet of poly-A-RNA was selected as this is the minimal size that let in experiments to consistent prion conversion, while photo-degradation below the five nucleotide threshold drastically lowered the rate of conversion. ^(36-37)^ Our docking protocol allows for free rotation around all single bonds in the RNA molecule. The ten highest-scoring docked systems for all three target structures were collected for the next stage of model generation. These complexes were examined for common regions of protein-RNA interaction and compared to the ones seen in our previous study. The D178N mutation 129M-178N has for all 10 complexes as most probable binding site the one which we called binding site 3 (residues 135-145) in our previous study. The M129V sequence change (129V-178D) results in eight complexes with binding site 2 (residues 111-121) and two complexes with binding site 1 (residues 21-31). The third system, characterized by residue changes M129V and D178N (129V-178N), has again only binding sites also seen in our previous study: six complexes with binding site 1, one complex with binding site 2 and three complexes with binding site 3. We then evaluated the stability of all thirty complexes in short (10 ns) long molecular dynamics simulations with a temperature of 310K and pressure of 1 bar, looking for RNA detachment from he protein. No such detachment was seen for the D178N mutant 129M-178N. For the 129V-178D system, the RNA did detach for two of the eight complexes with binding site 2. For the system with both residue alterations M129V and D178N (129V-178N), the RNA detached from the Prion not only in the sole complex with binding site 2 but also in one of the six complexes with binding site 1. Note that in our previous study of prion RNA interaction^(17)^, we encountered similar events of RNA detachment, all involving binding site 2. In the present study, we considered for long molecular dynamics runs only complexes where we did not observe RNA detachment in the short runs. Out of the remaining cluster we selected as start configuration for the long runs for each system and each observed binding site the one that had the lowest root-man-square-deviation (RMSD) in the helix A region at the end of the above described short runs. In this way we tried to minimize any possible bias in our complexes toward helix instability. **Table 1** presents a matrix of systems and binding sites with the number of structures and long molecular dynamics runs for each case. As a control, we also evaluate the stability of the isolated prions, by following molecular dynamics trajectories of same length and generated with the same simulation protocol, with the prion having the same start configuration as in the corresponding complex.

### Simulation Protocol

All our molecular dynamics simulations rely on the GROMACS software package version 4.6.5^(38)^, using the CHARMM36 force field with associated nucleic acid parameters ^39,46^ and TIP3P water model^(43-44)^ to model interactions between protein, RNA and water. The prion was put at the center cubic box with at least 12 Å distance between the boundary of the box and the protein-RNA complex, and he box filled with water molecules. Periodic boundary conditions are used and electrostatic interactions calculated via the PME algorithm.^45-46^ Due to the size of the system, and the potential for steric clashes during solvation, the solvated model was relaxed first by steepest descent energy minimization followed a 2 ns molecular dynamic simulation using NVT protocol and a subsequent 2ns simulation using NPT protocol.

For integration of the equations of motion a 2 fs time step is used, with hydrogen atoms constrained by the LINC algorithm^(47)^ and water constrained with the Settle algorithm^(48)^. The temperature of the system is kept at 310 K by the Parrinello-Donadio-Bussi thermostat^(48-49)^ (τ = 0.1 fs), and the pressure at 1 bar by h Parrinello-Rahman algorithm (τ = 1 fs).^(50)^. A group cutoff scheme was selected, with neighbor search using a grid-cutoff scheme with a cutoff distance of 1.5 nm. Electrostatic interactions outside these dimensions are handled by Particle Mesh Ewald with cubic interpolation and grid dimensions set to 0.15nm.

For each of he systems listed in **Table 1** we run three molecular dynamic trajectories differing in the initial velocity distributions to get a simple estimate of the statistical fluctuations between trajectories. Data are saved every 4ps for further analysis. Using the internal tools of GROMACS we measured: root-means-square deviations of the Cα atoms (RMSD), secondary structure contents, contact distances and hydrogen bonding. Configurations are visualized using PYMOL and VMD.

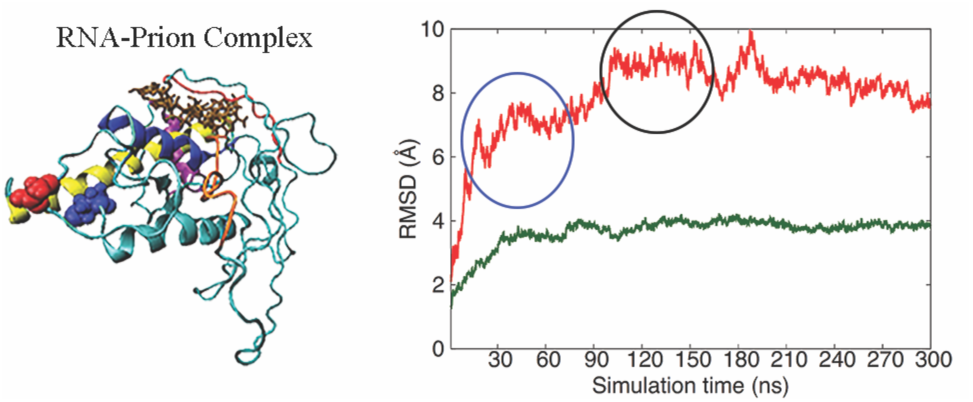
Table of Content Figure:

## ACKNOWLEDGMENT

We thank Jarek Meller (University of Cincinnati) for pointing us to the Familial Fatal Insomnia related prion sequences and for stimulating discussions during CBSB2017. The simulations in this work were done using XSEDE resources funded under project MCB160005 and on the BOOMER cluster of the University of Oklahoma. We acknowledge financial support from the National Science Foundation under grant IOS1453613 and the National Institutes of Health under grant GM120634.

## References

1. Deleault, N.-R.; Harris, B.-T.; Rees, J. R.; Supattapone, S. Formation of native prions from minimal components in vitro. Proc Natl Acad Sci U S A 2007, 104, 9741–6.

2. Westergard, L.; Christensen, H.-M.; Harris, D.-A. The cellular prion protein (PrP(C)): its physiological function and role in disease. Biochim Biophys Acta 2007, 1772, 629–644.

3. Bremer, J.; Baumann, F.; Tiberi, C.; Wessig, C.; Fischer, H.; Schwarz, P.; Steele, A.-D.; Toyka, K.-V.; Nave, K.-A.; Weis, J.; Aguzzi, A. Axonal prion protein is required for peripheral myelin maintenance. Nat Neurosci 2010, 13, 310–318.

4. Caiati, M.-D.; Safiulina, V.-F.; Fattorini, G.; Sivakumaran, S.; Legname, G.; Cherubini, E. PrPC controls via protein kinase A the direction of synaptic plasticity in the immature hippocampus. J Neurosci 2013, 33, 2973–2983.

5. Buhimschi, I.-A.; Nayeri, U.-A.; Zhao, G.; Shook, L.-L.; Pensalfini, A.; Funai, E.-F.; Bernstein, I.-M.; Glabe, C.-G.; Buhimschi, C.-S. Protein misfolding, congophilia, oligomerization, and defective amyloid processing in preeclampsia. Sci Transl Med 2014, 6, DOI: 10.1126/scitranslmed.3008808.

6. Kouza, M.; Banerji, A.; Kolinski, A.; Buhimschi, I.-A.; Kloczkowski, A. Oligomerization of FVFLM peptides and their ability to inhibit beta amyloid peptides aggregation: consideration as a possible model. Phys Chem Chem Phys 2017, 19, 2990–2999.

7. Pan, K.-M.; Baldwin, M.; Nguyen, J.; Gasset, M.; Serban, A.; Groth, D.; Mehlhorn, I.; Huang, Z.; Fletterick, R.-J.; Cohen, F.-E.; Prusiner, S.-B. Conversion of alpha-helices into betasheets features in the formation of the scrapie prion proteins. Proc Natl Acad Sci U S A 1993, 90, 10962–10966.

8. Norstrom, E.-M.; Mastrianni, J.-A., The charge structure of helix 1 in the prion protein regulates conversion to pathogenic PrPSc. J Virol 2006, 80, 8521–8529.

9. Singh, J.; Kumar, H.; Sabareesan, A.-T.; Udgaonkar, J.-B. Rational stabilization of helix 2 of the prion protein prevents its misfolding and oligomerization. J Am Chem Soc 2014, 136, 16704–16707.

10. Prusiner, S.-B. Prion diseases and the BSE crisis. Science 1997, 278, 245–251.

11. Singh, J.; Udgaonkar, J.-B. Dissection of conformational conversion events during prion amyloid fibril formation using hydrogen exchange and mass spectrometry. J Mol Biol 2013, 425, 3510–3521.

12. Schlepckow, K.; Schwalbe, H., Molecular mechanism of prion protein oligomerization at atomic resolution. Angew Chem Int Ed Engl 2013, 52, 10002–10005.

13. Tycko, R.; Savtchenko, R.; Ostapchenko, V.-G.; Makarava, N.; Baskakov, I.-V. The alpha-helical C-terminal domain of full-length recombinant PrP converts to an in-register parallel beta-sheet structure in PrP fibrils: evidence from solid state nuclear magnetic resonance. Biochemistry 2010, 49, 9488–9497.

14. Salamat, K.; Moudjou, M.; Chapuis, J.; Herzog, L.; Jaumain, E.; Beringue, V.; Rezaei, H.; Pastore, A.; Laude, H.; Dron, M. Integrity of helix 2-helix 3 domain of the PrP protein is not mandatory for prion replication. J Biol Chem 2012, 287, 18953–18964.

15. Prusiner, S.-B. Prions. Proc Natl Acad Sci U S A 1998, 95, 13363–13383.

16. Miller, M.-B.; Geoghegan, J.-C.; Supattapone, S. Dissociation of infectivity from seeding ability in prions with alternate docking mechanism. PLoS Pathog 2011, 7, DOI: 10.1371/journal.ppat.1002128

17. Alred, E.-J.; Nguyen, M.; Martin, M.; Hansmann, U.-H.-E. Molecular dynamics simulations of early steps in RNA-mediated conversion of prions. Protein Sci 2017, 26, 1524–1534.

18. Zurawel, A.-A.; Walsh, D.-J.; Fortier, S.-M.; Chidawanyika, T.; Sengupta, S.; Zilm, K.; Supattapone, S. Prion nucleation site unmasked by transient interaction with phospholipid cofactor. Biochemistry 2014, 53, 68–76.

19. Frau-Mendez, M.-A.; Fernandez-Vega, I.; Ansoleaga, B.; Blanco Tech, R.; Carmona Tech, M.; Antonio Del Rio, J.; Zerr, I.; Llorens, F.; Jose Zarranz, J.; Ferrer, I. Fatal familial insomnia: mitochondrial and protein synthesis machinery decline in the mediodorsal thalamus. Brain Pathol 2017, 27 (1), 95–106.

20. Saa, P.; Harris, D.-A.; Cervenakova, L. Mechanisms of prion-induced neurodegeneration. Expert Rev Mol Med 2016, 18, DOI: 10.1017/erm.2016.8.

21. Forloni, G.; Tettamanti, M.; Lucca, U.; Albanese, Y.; Quaglio, E.; Chiesa, R.; Erbetta, A.; Villani, F.; Redaelli, V.; Tagliavini, F.; Artuso, V.; Roiter, I. Preventive study in subjects at risk of fatal familial insomnia: Innovative approach to rare diseases. Prion 2015, 9, 75–89.

22. Synofzik, M.; Bauer, P.; Schols, L. Prion mutation D178N with highly variable disease onset and phenotype. J Neurol Neurosurg Psychiatry 2009, 80, 345–346.

23. Golanska, E.; Sieruta, M.; Corder, E.; Gresner, S.-M.; Pfeffer, A.; Chodakowska-Zebrowska, M.; Sobow, T.-M.; Klich, I.; Mossakowska, M.; Szybinska, A.; Barcikowska, M.; Liberski, P.-P. The prion protein M129V polymorphism: longevity and cognitive impairment among Polish centenarians. Prion 2013, 7, 244–247.

24. Laplanche, J.-L.; Delasnerie-Laupretre, N.; Brandel, J.-P.; Chatelain, J.; Beaudry, P.; Alperovitch, A.; Launay, J.-M. Molecular genetics of prion diseases in France. French Research Group on Epidemiology of Human Spongiform Encephalopathies. Neurology 1994, 44 (12), 2347–2351.

25. Webb, B.; Sali, A. Comparative Protein Structure Modeling Using MODELLER. Curr Protoc Bioinformatics 2014, 47, 1–32.

26. Marti-Renom, M.-A.; Stuart, A.-C.; Fiser, A.; Sanchez, R.; Melo, F.; Sali, A. Comparative protein structure modeling of genes and genomes. Annu Rev Biophys Biomol Struct 2000, 29, 291–325.

27. Yang, J.; Yan, R.; Roy, A.; Xu, D.; Poisson, J.; Zhang, Y. The I-TASSER Suite: protein structure and function prediction. Nat Methods 2015, 12, 7–8.

28. Roy, A.; Kucukural, A.; Zhang, Y. I-TASSER: a unified platform for automated protein structure and function prediction. Nat Protoc 2010, 5, 725–738.

29. Zhang, Y. I-TASSER server for protein 3D structure prediction. BMC Bioinformatics 2008, 9, 40.

30. Colovos, C.; Yeates, T.-O. Verification of protein structures: patterns of nonbonded atomic interactions. Protein Sci 1993, 2, 1511–1519.

31. Lovell, S.-C.; Davis, I. W.; Arendall, W.-B.; de Bakker, P.-I.; Word, J.-M.; Prisant, M. G.; Richardson, J.-S.; Richardson, D.-C. Structure validation by Calpha geometry: phi,psi and Cbeta deviation. Proteins 2003, 50, 437–450.

32. Wallner, B.; Elofsson, A. Can correct protein models be identified? Protein Sci 2003, 12 (5), 1073–1086.

33. Maiti, R.; Van Domselaar, G.-H.; Zhang, H.; Wishart, D.-S. SuperPose: a simple server for sophisticated structural superposition. Nucleic Acids Res 2004, 32, 590–594.

34. Peltonen, K.; Colis, L.; Liu, H.; Trivedi, R.; Moubarek, M.-S.; Moore, H.-M.; Bai, B.; Rudek, M.-A.; Bieberich, C.-J.; Laiho, M. A targeting modality for destruction of RNA polymerase I that possesses anticancer activity. Cancer Cell 2014, 25, 77–90.

35. Hosen, M.-J.; Zubaer, A.; Thapa, S.; Khadka, B.; De Paepe, A.; Vanakker, O.-M. Molecular docking simulations provide insights in the substrate binding sites and possible substrates of the ABCC6 transporter. PLoS One 2014, 9, DOI: 10.1371/journal.pone.0102779.

36. Piro, J.-R.; Supattapone, S. Photodegradation illuminates the role of polyanions in prion infectivity. Prion 2011, 5, 49–51.

37. Piro, J.-R.; Harris, B.-T.; Supattapone, S. In situ photodegradation of incorporated polyanion does not alter prion infectivity. PLoS Pathog 2011, r7, DOI: 10.1371/journal.ppat.1002001.

38. Pronk, S.; Pall, S.; Schulz, R.; Larsson, P.; Bjelkmar, P.; Apostolov, R.; Shirts, M.-R.; Smith, J.-C.; Kasson, P.-M.; van der Spoel, D.; Hess, B.; Lindahl, E. GROMACS 4.5: a High-Throughput and Highly Parallel Open Source Molecular Simulation Toolkit. Bioinformatics 2013, 29, 845–854.

39. MacKerell, A.-D.; Bashford, D.; Bellott, M.; Dunbrack, R.-L.; Evanseck, J.-D.; Field, M.-J.; Fischer, S.; Gao, J.; Guo, H.; Ha, S.; Joseph-McCarthy, D.; Kuchnir, L.; Kuczera, K.; Lau, F.-T.; Mattos, C.; Michnick, S.; Ngo, T.; Nguyen, D.-T.; Prodhom, B.; Reiher, W.-E.; Roux, B.; Schlenkrich, M.; Smith, J.-C.; Stote, R.; Straub, J.; Watanabe, M.; Wiorkiewicz-Kuczera, J.; Yin, D.; Karplus, M. All-atom empirical potential for molecular modeling and dynamics studies of proteins. J Phys Chem B 1998, 102, 3586–3616.

40. Mackerell, A.-D.; Feig, M.; Brooks, C.-L. Extending the treatment of backbone energetics in protein force fields: limitations of gas-phase quantum mechanics in reproducing protein conformational distributions in molecular dynamics simulations. J Comput Chem 2004, 25, 1400–1415.

41. Denning, E.-J.; Priyakumar, U.-D.; Nilsson, L.; Mackerell, A.-D. Impact of 2’-hydroxyl sampling on the conformational properties of RNA: update of the CHARMM all-atom additive force field for RNA. J Comput Chem 2011, 32, 1929–1943.

42. Best, R.-B.; Zhu, X.; Shim, J.; Lopes, P.-E.; Mittal, J.; Feig, M.; Mackerell, A.-D. Optimization of the additive CHARMM all-atom protein force field targeting improved sampling of the backbone phi, psi and side-chain chi(1) and chi(2) dihedral angles. J Chem Theory Comput 2012, 8, 3257–3273.

43. Mahoney, M.-W.; Jorgensen, W.-L. A five-site model for liquid water and the reproduction of the density anomaly by rigid, nonpolarizable potential functions. The Journal of Chemical Physics 2000, 112, 8910–8922.

44. Jorgensen, W.-L.; Chandrasekhar, J.; Madura, J.-D.; Impey, R.-W.; Klein, M.-L. Comparison of simple potential functions for simulating liquid water. J. Chem. Phys. 1983, 79, 926–935.

45. Darden, T.; York, D.; Pedersen, L. Particle mesh Ewald: An N log(N) method for Ewald sums in large systems. The Journal of Chemical Physics 1993, 98, 10089–10092.

46. Essmann, U.; Perera, L.; Berkowitz, M.-L.; Darden, T.; Lee, H.; Pedersen, L.-G. A smooth particle mesh Ewald method. The Journal of Chemical Physics 1995, 103, 8577–8593.

47. Hess, B., P-LINCS: A parallel linear constraint solver for molecular simulation. J. Chem. Theory Comput. 2008, 4, 116–122.

48. Miyamoto, S.; Kollman, P.-A. Settle - an analytical version of the shake and rattle algorithm for rigid water models. J. Comput. Chem. 1992, 13, 952–962.

49. Bussi, G.; Donadio, D.; Parrinello, M. Canonical sampling through velocity rescaling. J. Chem. Phys. 2007, 126, DOI: 10.1063/1.2408420.

50. Parrinello, M.; Rahman, A. Polymorphic transitions in single crystals: A new molecular dynamics method. J. Appl. Phys. 1981, 52, 7182–7190.

51. The PyMOL Molecular Graphics System, Version 1.8 Schrödinger, LLC.

